# Inhibiting NINJ1-dependent plasma membrane rupture protects against inflammasome-induced blood coagulation and inflammation

**DOI:** 10.1101/2023.08.30.555561

**Authors:** Jian Cui, Hua Li, Dien Ye, Guoying Zhang, Yan Zhang, Ling Yang, Martha M.S. Sim, Jeremy P. Wood, Yinan Wei, Zhenyu Li, Congqing Wu

## Abstract

Systemic blood coagulation accompanies inflammation during severe infection like sepsis and COVID. We’ve previously established a link between pyroptosis, a vital defense mechanism against infection, and coagulopathy. During pyroptosis, the formation of gasdermin-D (GSDMD) pores on the plasma membrane leads to the release of tissue factor (TF)-positive microvesicles (MVs) that are procoagulant. Mice lacking GSDMD release fewer TF MVs. However, the specific mechanisms leading from activation of GSDMD to MV release remain unclear. Plasma membrane rupture (PMR) in pyroptosis was recently reported to be actively mediated by the transmembrane protein Ninjurin-1 (NINJ1). Here we show that NINJ1 promotes procoagulant MV release during pyroptosis. Haploinsuffciency or glycine inhibition of NINJ1 limited the release of procoagulant MVs and inflammatory cytokines and partially protected against blood coagulation and lethality triggered by bacterial flagellin. Our findings suggest a crucial role for NINJ1-dependent PMR in inflammasome-induced blood coagulation and inflammation.

## Introduction

Inflammasomes are complex intracellular sensors that detect pathogens by sensing pathogen-associated molecular patterns (PAMPs) (*1, 2*). The activation of inflammasome and subsequent pyroptosis is critical in host defense against pathogens (*3–5*). Pyroptosis exposes intracellular pathogens through plasma membrane rupture (PMR) (*6*). However, overactive inflammasomes, common in conditions like sepsis and COVID (*7–11*), can trigger systemic blood coagulation and tissue and organ dysfunction (*12–16*).

Tissue factor (TF) is a membrane protein and the primary initiator of the coagulation cascade, essential for hemostasis to stop bleeding upon injury (*17, 18*). Increased TF activity is associated with consumption coagulopathy linked to sepsis and COVID (*19–22*). Our previous study has shown that TF-positive MVs released by pyroptotic macrophages trigger systemic coagulation and lethality in experimental sepsis (*23*). However, the molecular mechanisms by which pyroptosis releases procoagulant MVs remain elusive. Pyroptotic cells undergo PMR at the late stage when the cell membrane is beyond repair, where the direct blebbing and pinching of the plasma membrane releases MVs to remove membrane damaged with GSDMD pores (*24–28*). PMR could accelerate MV release as the entire plasma membrane is subject to dissolve as MVs. Kayagaki et al. recently reported that Ninjurin-1 (NINJ1) oligomerization activates PMR, a process formerly thought to be passive, during lytic cell death like necrosis and pyroptosis (*29*). The role of NINJ1 in releasing procoagulant MVs and initiating systemic coagulation after inflammasome activation remains unknown.

In this study, we show that *Ninj1* haploinsufficiency inhibits the release of TF-positive MVs and provides protection against systemic coagulation. Reduced NINJ1 also limits the release of inflammatory cytokines. We find that a single dose of glycine injection has similar protective effects as they prevent PMR. Our findings underscore the essential role of NINJ1-dependent PMR in inflammasome-induced blood coagulation, thus expanding our understanding of the dual role that inflammasomes play in host defense.

## Results and discussion

### *Ninj1* haploinsufficiency protects against flagellin-induced blood coagulation

We aimed to understand the role of NINJ1 on inflammasome-induced blood coagulation, using heterozygous *Ninj1^+/−^* mice with decreased NINJ1 protein abundance (Fig S1). This approach was justified by our observation that total NINJ1 deficiency could lead to partial embryonic lethality and the reported occurrence of hydrocephalus in surviving mice (*30*).

To induce blood coagulation and lethality, we applied our established protocol the involves injection of mice with purified bacterial flagellin, fused with the cytosolic translocation domain of anthrax lethal factor (LFn). In the presence of anthrax protein protective agent (PA), this flagellin fusion protein (LFn-Fla) can be transported into the cytoplasm (*4, 31*). Flagellin is a PAMP known to activate the NAIP/NLRC4 inflammasome and trigger pyroptosis (*3, 4, 32, 33*). We confirmed robust inflammasome activation and pyroptosis in mouse primary bone marrow-derived macrophages (BMDMs) upon stimulating with purified Fla (LFn-Fla plus PA) (Fig S2A-B).

Our earlier work has shown that inflammasome-induced systemic coagulation is marked by prolonged prothrombin time (PT) and increased plasma thrombin-antithrombin (TAT) (*23*), frequently observed in patients with disseminated intravascular coagulation (DIC) and COVID (*12–16, 34*). As expected, intravenous injection of Fla activated systemic coagulation in wild-type *Ninj1^+/+^* mice, demonstrated by a significant increase of PT and a rise in plasma TAT (Fig 1A-B). Notably, when challenged with Fla, heterozygous *Ninj1^+/−^* mice showed lower PT and almost complete suppression of the elevation in plasma TAT compared with their wild-type littermates (Fig 1A-B). Our results suggest that a single copy deletion of *Ninj1* is adequate for significant inhibition of inflammasome-induced blood coagulation.

**Figure 1.**
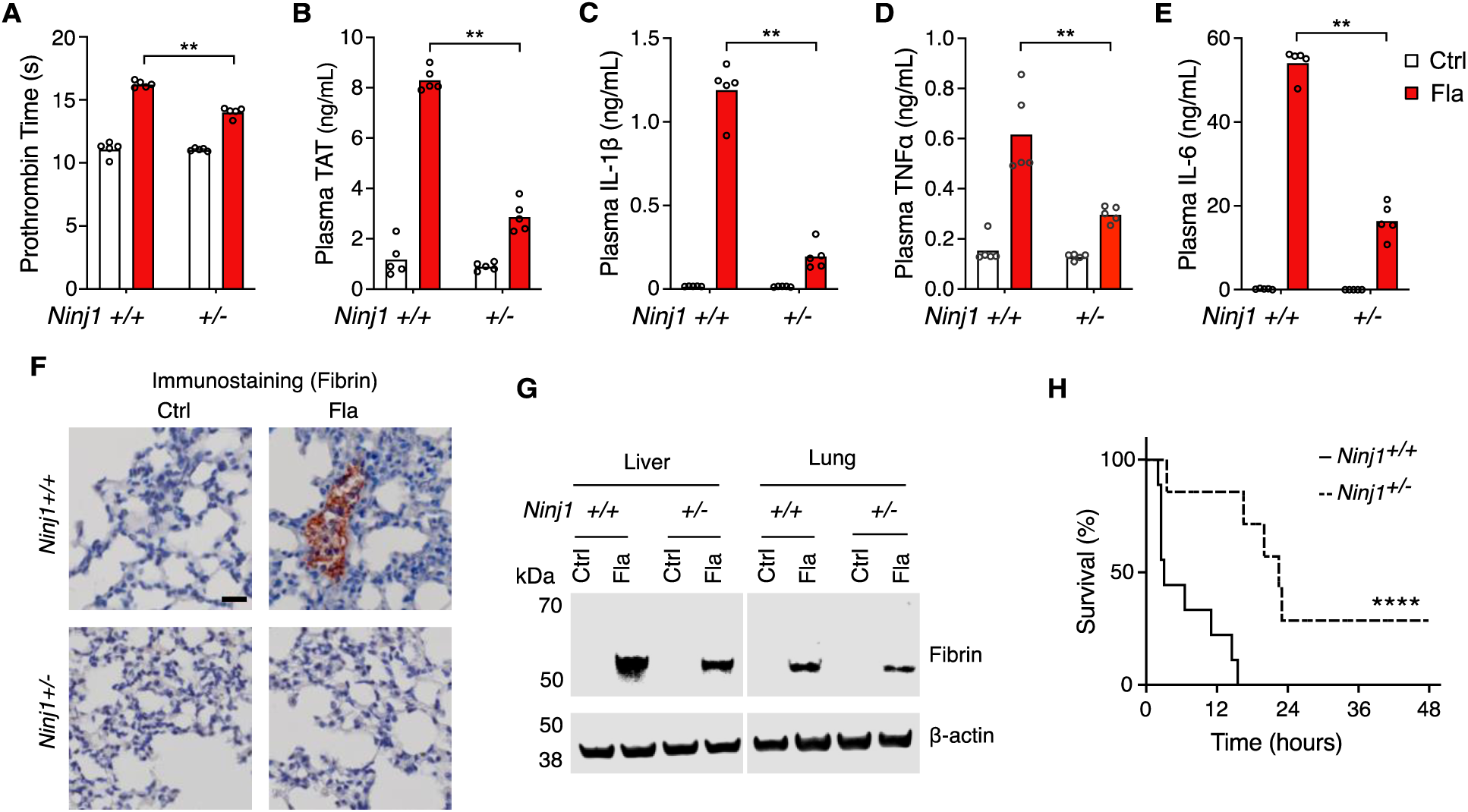
NINJ1 is critical for flagellin-induced systemic coagulation, inflammation, and lethality. (**A–E**) Mice were injected intravenously with Ctrl (saline) or Fla (500 ng LFn-Fla plus 3 μg PA). Blood was collected 90 minutes after Ctrl or Fla injection. Prothrombin time (**A**), plasma TAT (**B**), and plasma cytokines (**C-E**) were measured. Circles represent individual mouse, with bars donating mean. ** p < 0.01 (two-way ANOVA with Holm-Sidak multiple comparisons). (**F-G**) Mice were injected intravenously with Ctrl or Fla. After 90 minutes, mice were euthanized and perfused with PBS, and tissues were isolated. (**F**) Lung sections were stained with the anti-fibrin monoclonal antibody (59D8). Scale bar denotes 20 μm. Data are representative of 3 independent experiments (biological replicates). (**G**) Fibrin in the liver and lungs was detected by immunoblot with the anti-fibrin monoclonal antibody (59D8). Data are representative of 3 independent experiments (biological replicates). (**H**) Mice were injected intravenously with a lethal dose of Fla (2.5 μg LFn-Fla plus 6 μg PA). Kaplan-Meier survival plots for mice challenged with Fla are shown. n = 7-9. **** p < 0.0001 versus WT (log rank test [Mantel-Cox]).

We also found that *Ninj1* haploinsufficiency hindered the release of cytokines, as shown by decreased plasma concentrations of cytokines like IL-1β, TNFα, and IL-6 (Fig 1C-E). The decrease in cytokine concentrations could be attributed to limited release of danger-associated molecular patterns (DAMPs) due to less effective PMR in *Ninj1^+/−^*mice. It’s worth noting that GSDMD facilitates IL-1β and NINJ1 deficiency does not affect pyroptosis-induced IL-1β release into cell culture supernatant from BMDMs (ref). Given that *Ninj1* haploinsufficiency did not change GSDMD expression (Fig 1S), there might be unrecognized mechanisms in pyroptosis-induced IL-1β *in vivo*.

While our earlier research suggested that cell death, not the release of inflammatory cytokines like IL-1β, is crucial for the over-activation of blood coagulation triggered by inflammasomes (*23*), these cytokines are vital for maintaining inflammation. Our findings indicate their release is facilitated by NINJ1.

### Fibrin deposition and lethality resulting from inflammasome activation depend on NINJ1

Fibrin deposition in tissues can restrict blood supply, potentially resulting in tissue and organ damage and even death (*12*). Following the Fla challenge, we noted fibrin deposition in the lungs of wild-type mice through immunostaining with a fibrin-specific antibody 59D8 (ref) (Fig 1F), which was consistent with H&E staining (Fig 3S). However, no signs of fibrin deposition were observed in lung sections of *Ninj1^+/−^* mice, likely due to the technical limitations of immunostaining on thin (5-μm) tissue sections, compounded by the potential for uneven fibrin distribution. By using immunoblot to examine a homogenized whole lobe of the lung or liver, we detected fibrin deposition in the liver and lungs of *Ninj1^+/−^*mice, but to a much less extent compared with their *Ninj1^+/+^* littermates (Fig 1G). Furthermore, *Ninj1* haploinsufficiency partially rescued flagellin-induced lethality (Fig 1H). These findings underscore the significant role of NINJ1 protein in inflammasome-triggered fibrin deposition in tissues and lethality.

### NINJ1 plays a critical role in systemic coagulation in response to *E. coli* infection

Systemic blood coagulation is a common complication of pathogen-triggered sepsis (*35, 36*). To investigate the role of NINJ1 in bacterial infection-induced blood coagulation, *Ninj1^+/−^* mice were infected with cultured *E. coli* at a dose of 2 x 10^8^ pfu per mouse. Consistently with the findings from the Fla challenge, *E. coli* infection-induced blood coagulation was limited by *Ninj1* haploinsufficiency. As shown in Fig 2A-B, *Ninj1^+/−^* mice exhibited a lower PT and decreased plasma TAT following *E. coli* infection. Furthermore, *Ninj1^+/−^*mice displayed alleviated cytokine release in response to *E. coli* infection (Fig 2C-E). Collectively, these data demonstrate that reduced expression of NINJ1 confers protection against *E. coli* infection-induced blood coagulation. In addition, the alleviation of cytokine release in *Ninj1^+/−^*mice supports the role of NINJ1 in modulating the inflammatory response to *E. coli* infection. These findings highlight a potential role of NINJ1 in the pathogenesis of bacterial infection, including blood coagulation and the cytokine storm.

**Figure 2.**
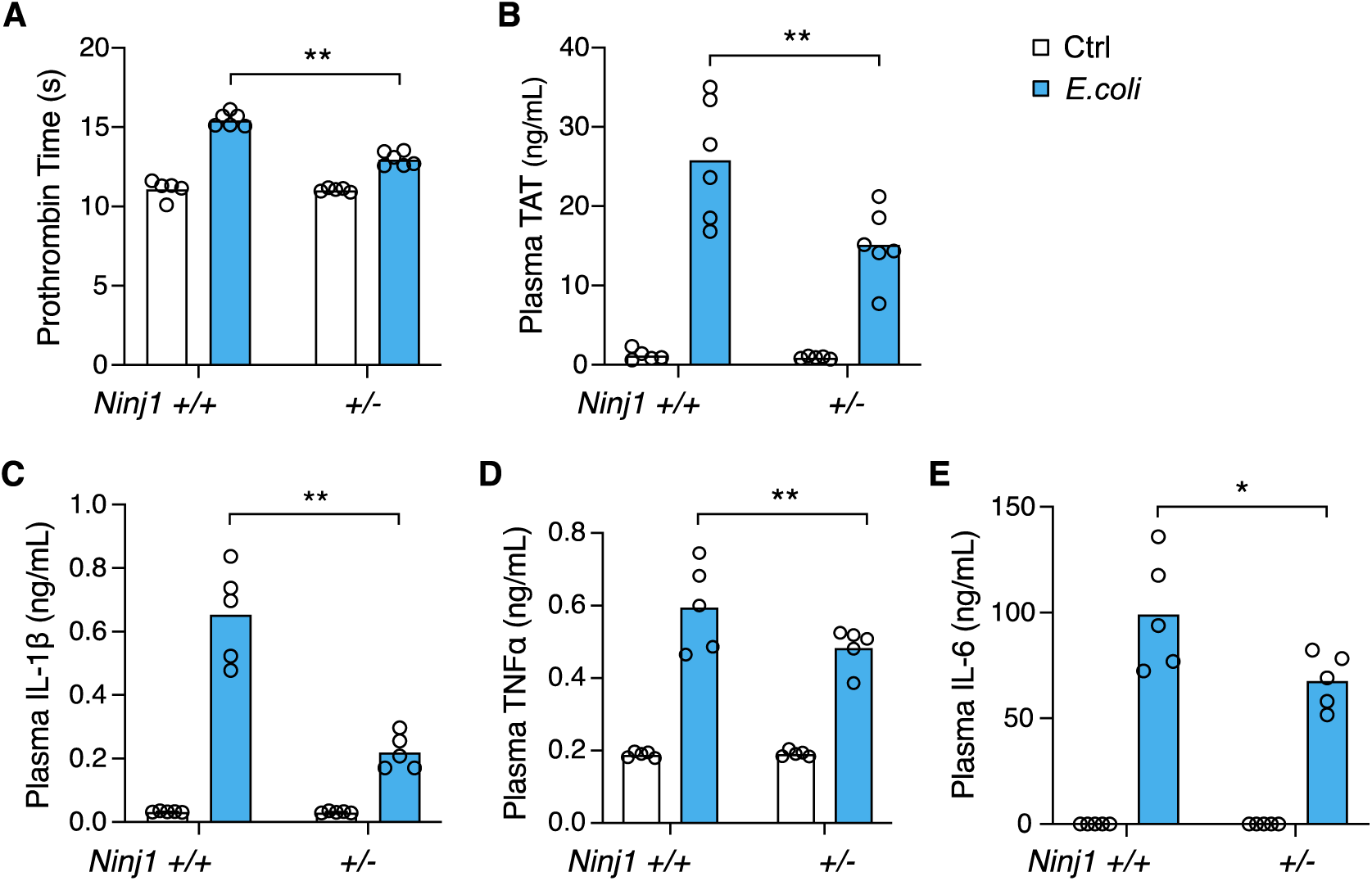
*E. coli* infection-induced blood coagulation is limited in *Ninj*1^+/−^ mice. (**A–E**) Mice were injected intravenously with Ctrl (saline) or *E. coli* (2 x 10^8^ cfu per mouse). Blood was collected 6 hours afterwards. Prothrombin time (**A**), plasma TAT (**B**), and plasma cytokines (**C-E**) were measured. Circles represent individual mouse, with bars donating mean. * p < 0.05, ** p < 0.01 (two-way ANOVA with Holm-Sidak multiple comparisons).

**Figure 3.**
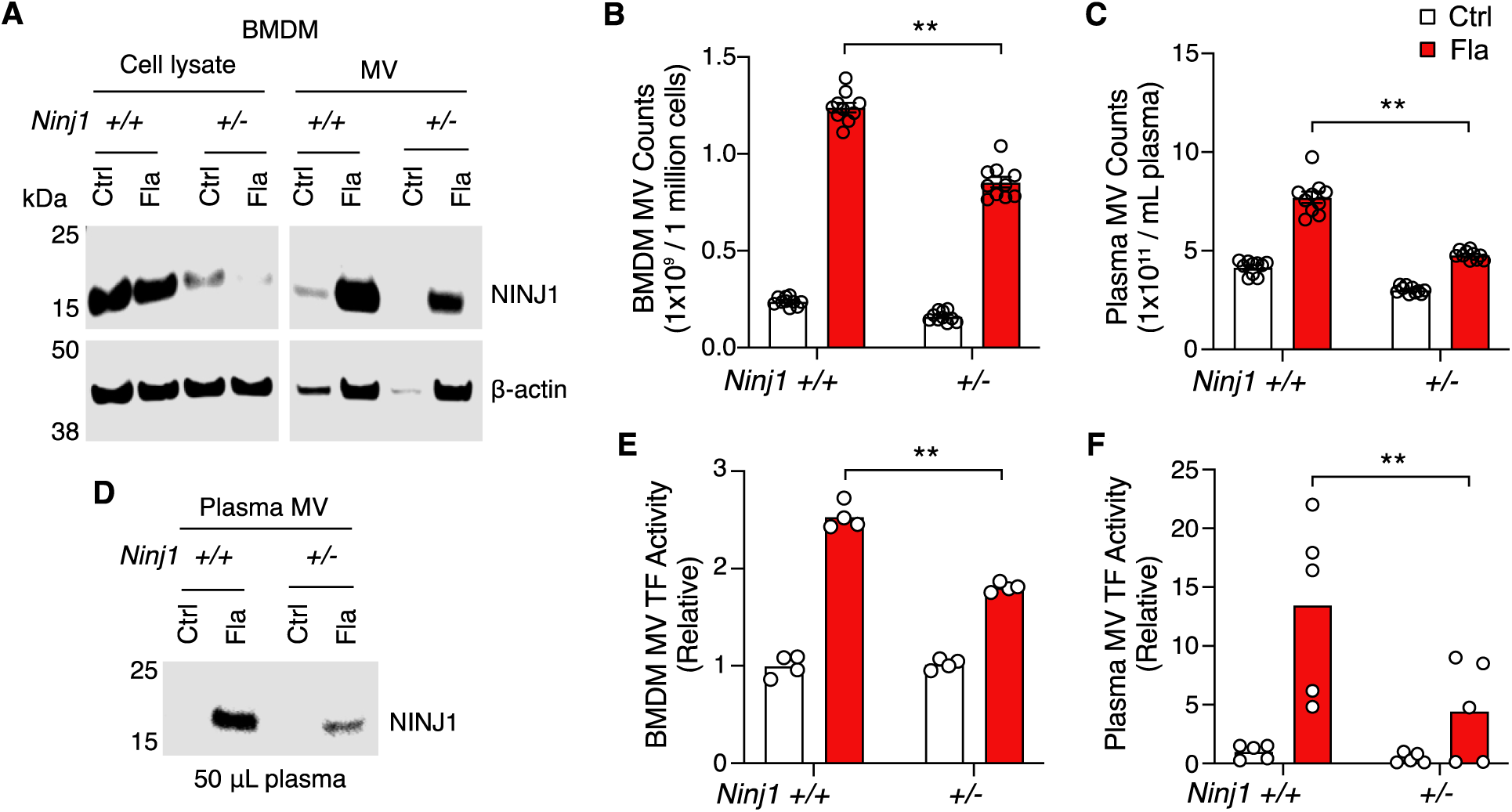
PMR promotes the release of procoagulant MVs. (**A-B, E**) BMDMs were incubated with Ctrl (PBS) or Fla (1 μg/mL LFn-Fla plus 1 μg/mL PA). Cell culture supernatant and MVs were collected after 90 minutes of incubation. (**A**) NINJ1 in the cell lysate and MVs was detected by immunoblot. (**B**) BMDM MVs were counted with Nanosight. (**E**) BMDM MV TF activity. Circles represent individual mouse, with bars donating mean. ** p < 0.01 (two-way ANOVA with Holm-Sidak multiple comparisons). (**C, D, F**) Mice were injected intravenously with Ctrl (saline) or Fla (500 ng LFn-Fla plus 3 μg PA). Blood was collected 90 minutes after Ctrl or Fla injection. (**C**) Plasma MVs were counted with NanoSight. (**D**) NINJ1 in plasma MVs was detected with equal plasma volumes by immunoblot. (**F**) Plasma MV TF activity. Circles represent individual mouse, with bars donating mean. ** p < 0.01 (two-way ANOVA with Holm-Sidak multiple comparisons).

### NINJ1-dependent PMR promotes procoagulant MV release

In our previous work, we established that macrophage pyroptosis promotes the release of procoagulant MVs (*23*). Yet, the cellular processes linking pyroptosis to MV release remain to be defined. We hypothesize that PMR during pyroptosis could be the molecular event responsible for the release of these MVs. To test this hypothesis, we used BMDMs isolated from *Ninj1^+/+^* and *Ninj1^+/−^*littermates. We observed a significant reduction in PMR, indicated by the decreased lactate dehydrogenase (LDH) in culture medium, in *Ninj1^+/−^* cells (Fig S3A), while cell death, measured by ATP levels, remained unaffected (Fig S3B). This is consistent with previous studies suggesting that NINJ1 mediates PMR but not cell death (*29, 37*).

Notably, we detected NINJ1 protein in MVs released from *Ninj1^+/+^*BMDMs (Fig 3A). The protein level was decreased in MVs derived from *Ninj1^+/−^* BMDMs (Fig 3A), likely due to reduced MV release and reduced NINJ1 protein in *Ninj1^+/−^* cells (Fig 3B). Consistently, *in vivo*, NINJ1 was detected in plasma MVs from *Ninj1^+/+^*mice exposed to Fla (Fig 3D). *Ninj1^+/−^*mice also had decreased amount of NINJ1 from MVs, likely due to fewer plasma MVs (Fig 3C). The reduced quantity of MVs from both plasma and BMDMs in *Ninj1^+/−^* mice correlated with lower TF activity (Fig 3E and 3F). Our data illustrate that a decline in NINJ1 expression significantly reduces PMR and MV release. These findings provide compelling evidence supporting a critical role of NINJ1-dependent PMR in procoagulant MV release. Furthermore, MV shedding removes NINJ1 from the plasma membrane, hinting at a possible role of NINJ1 in cell repair.

### Glycine inhibition of NINJ1 alleviates pyroptosis-induced blood coagulation

It has long been recognized that glycine buffering protects against PMR during cell death, such as pyroptosis and necrosis (*38, 39*). The exact mechanisms underlying the glycine protective effects remained unclear until Borges et al. revealed that glycine inhibits NINJ1 oligomerization and shields cells from rupture (*37*). Yet, whether glycine buffering would block the release of PMR-induced TF MVs or limit pyroptosis-induced coagulation *in vivo* is unknown. To complement our genetic approach, we injected a glycine solution into wild-type mice, prior to Fla injection, aiming to inhibit NINJ1.

Intriguingly, following the Fla challenge, glycine injection led to a notable decrease in PT and plasma TAT (Fig 4A and 4B), demonstrating significant protection against flagellin-induced blood coagulation. Consistently, mice treated with glycine exhibited lower plasma MV TF activity post the Fla challenge (Fig 4C). *In vitro*, glycine buffering on isolated BMDMs protected against PMR, as denoted by a reduction LDH release (Fig S4A), and decreased MV TF activity (Fig S4B). However, we observed a discrepancy in MV counts between the *in vitro* and *in vivo* studies. Contrary to isolated BMDMs (Fig S4C), glycine-treated mice had higher MV counts irrespective of the Fla challenge (Fig 4D). We speculate that glycine might enhance MV release from cell types other than macrophages, independently of inflammasome activation. Lastly, glycine administration reduced the release of proinflammatory cytokines IL-1b and TNFa (consistent with previous research (*40*)), but not IL-6 (Fig 4E-G). In conclusion, the cytoprotective effect of glycine buffering resulted in improved outcomes in inflammasome-induced blood coagulation and inflammation.

**Figure 4.**
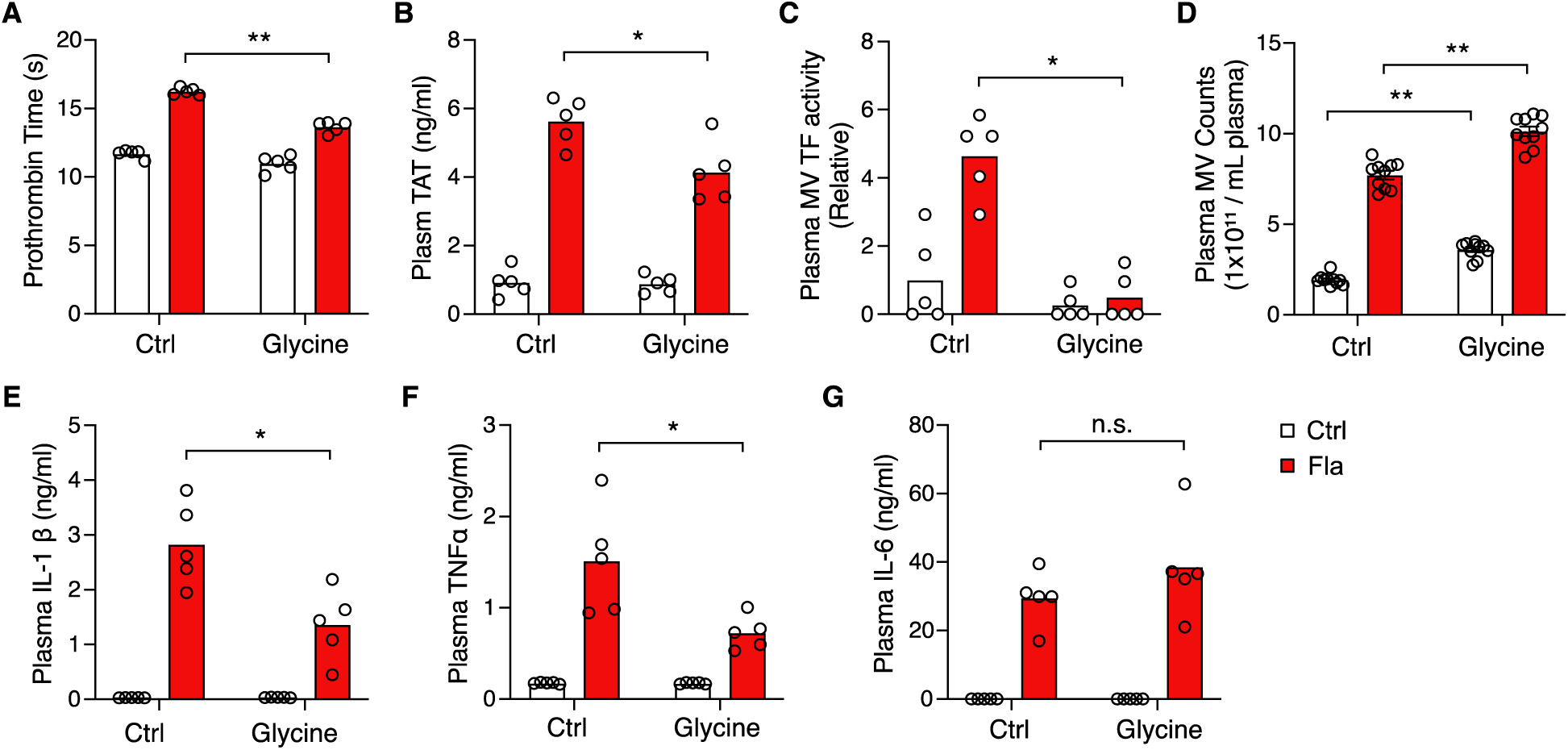
Glycine inhibition of NINJ1 blocks pyroptosis-induced blood coagulation. Mice were injected intravenously 50 μl of 0.5 M glycine 2 hours before administrating Ctrl (saline) or Fla (500 ng LFn-Fla plus 3 μg PA). Blood was collected 90 minutes afterwards. Prothrombin time (**A**), plasma TAT (**B**), plasma TF MV activity (**C**), plasma MV counts (D), and plasma cytokines (**E-G**) were measured. Circles represent individual mouse, with bars donating mean. * p < 0.05, ** p < 0.01, n.s. donates not significant (two-way ANOVA with Holm-Sidak multiple comparisons).

In summary, our study employed both genetic and pharmacological approaches to inhibit NINJ1-dependent PMR. Our findings suggest that PMR serves as a pivotal molecular event responsible for the release of procoagulant MVs and the induction of systemic coagulation and inflammation in an experimental model of sepsis. This novel mechanism sheds light on the potential for developing NINJ1 inhibition as a promising therapeutic strategy for alleviating acute inflammation and coagulation-associated organs and tissue and organ damage.

## Methods

### Mice

C57BL/6J, *Ninj1^+/+^,* and *Ninj1^+/−^* mice were housed in the University of Kentucky Animal Care Facility, following institutional and National Institutes of Health guidelines after approval by the Institutional Animal Care and Use Committee.

### Recombinant protein purification

Recombinant proteins were expressed using LPS free *E. coli* strain ClearColi BL21(DE3) (Lucigen Corporation, Cat#60810) at 37 °C for 4 hours with 500 μM IPTG after OD600 reached 0.6-0.8. Bacteria were collected and lysed in 50 mM Tris-HCL and 300 mM NaCl. Proteins containing a His-tag were purified by affinity chromatography using HisPur Ni-NTA resin (Thermo Fisher Scientific, Cat#88222). Proteins were then eluted with 250 mM imidazole in 50 mM Tris-HCl and 300 mM NaCl, and subsequently dialyzed against PBS to remove imidazole. Proteins concentrations were determined by measuring A280 before sterile filtration.

### *In vivo* Challenges

For flagellin challenge, purified PA and LFn-Fla in saline were administered via retro-orbital injection. Blood samples were collected at 90 minutes or 6 hours post-injection, followed by saline perfusion. Subsequently, tissue samples were harvested. For *E. coli* challenge, mice were injected intraperitoneally with *E. coli* saline solution. Blood samples were collected at 6 hours post-injection. For glycine *in vivo* experiment, mice were injected intravenously with glycine saline solution 2 hours before flagellin challenge. Blood samples were collected at 90 minutes after flagellin injection.

### BMDM isolation and differentiation

Mouse femur and tibia from one leg are harvested and rinsed in ice-cold PBS, followed by a brief rinse in 70% ethanol for 10-15 seconds. Both ends of the bones are then cut open, and the bone marrow is flushed out using a 10 ml syringe with a 26-gauge needle. The marrow is passed through a 19-gauge needle once to disperse the cells. After filtering through a 70-μm cell strainer, the cells are collected by centrifugation at 250 g for 5 minutes at 4 °C, then suspended in two 150 mm petri dish, each containing 25 ml of L-cell conditioned medium (RPMI-1640 supplemented with 10% FBS, 2mM L-Glutamine, 10mM HEPES, 15% LCM, and penicillin/streptomycin). After 3 days, 15 mL of LCM medium is added to each dish cells. The cells typically reach full confluency by days 5-7.

### BMDM culture and stimulation

BMDMs seeded into 12-well cell culture plate or 96-well cell culture plate at a density of 1 x 10^6^ cells/mL of RPMI-1640 medium (Thermo Fisher Scientific, Cat# 61870036) containing 10% of Fetal Bovine Serum (FBS) (Thermo Fisher Scientific, Cat# A3160502). BMDMs were allowed to settle overnight and refreshed with Opti-MEM (Thermo Fisher Scientific, Cat# 51985034) before purified proteins were added. To measure cytotoxicity, cell viability and MVs, supernatant was collected 90 minutes after stimulation.

### Cell viability and cytotoxicity

Cell viability of BMDMs was determined using CellTiter-Glo Luminescent Cell Viability Assay (Promega, Cat#G7572). Luminescence was recorded as an indicator of ATP concentrations in metabolically active cells. BMDMs cell cytotoxicity, as measured by LDH in the cell culture supernatant, was determined using CytoTox 96 Non-Radioactive Cytotoxicity Assay (Promega, Cat#G1780) according to manufacturer’s instruction.

### Prothrombin Time (PT)

Blood samples were collected from ketamine/xylazine-anaesthetized mice by cardiac puncture with a 23-gauge needle attached to a syringe pre-filled with 3.8% trisodium citrate as anticoagulant (final ratio at 1:10). Blood was centrifuged at 2,000 g for 20 minutes at 4 °C to obtain plasma. Prothrombin time (PT) was determined with Thromboplastin-D (Pacific Hemostasis, Cat#100357) in a manual setting according to manufacturer’s instruction, using CHRONO-LOG #367 plastic cuvette.

### Plasma TAT

Plasma was collected as mentioned above in PT. Plasma TAT concentrations were determined using a mouse TAT ELISA kit (Abcam, Cat#ab137994) at 1:50 dilution according to manufacturer’s instruction.

### ELISA

IL-1β, IL-6, and TNFα levels in culture supernatant and plasma were measured with ELISA kits from Thermo Fisher Scientific according to manufacturer’s instruction. Plates were read on a Cytation 5 at 450 and 570 nm.

### Tissue preparation and immunohistochemistry

Mice were perfused via both right and left ventricles with saline. Tissues were collected and embedded in paraffin, then sectioned at 5 mm. Anti-fibrin antibody 59D8 (kindly provided by Dr. Hartmut Weiler at Medical College of Wisconsin and Dr. Rodney M. Camire at the University of Pennsylvania) at 4 mg/ml was used for staining fibrin deposition, with M.O.M.® (Mouse on Mouse) ImmPRESS® HRP (Peroxidase) Polymer Kit (Vector, Cat#MP-2400) from vector laboratories according to manufacturer’s instruction for developing positive staining.

### Fibrin extraction for immunoblot

Frozen tissues were homogenized in 20 volumes (mg: μL) of T-PER tissue protein extraction reagent (Thermo Fisher Scientific, Cat#78510) containing cocktail inhibitor (Sigma, Cat#P8340). After centrifugation at 10,000 g for 10 minutes, supernatant was collected for β-actin detection. Pellets were then homogenized in 3 M urea and vortexed for 2 hours at 37 °C. After centrifugation at 14,000 g for 15 minutes, pellets were suspended in LDS sample buffer and vortexed at 65 °C for 30 minutes and ready for fibrin detection.

### Fluorescent immunoblot

For detection of Caspase-1, IL-1β, GSDMD, and NINJ1 by immunoblot, cells were washed with cold PBS and lysed with LDS sample buffer. Culture supernatant was precipitated with 1/10 volume of 2% sodium cholate and 1/10 volume of 100% trichloroacetic acid (TCA), and then dissolved in LDS sample buffer. TF was detected using anti-tissue factor (R&D, Cat#AF3178-SP) at 1:1000 dilution. Caspase-1 was detected using anti-Caspase-1 (Adipogen, Cat#AG-20B-0042-C100) at 1:1000 dilution. IL-1β was detected using anti-IL-1β (GeneTex, Cat#GTX74034) at 1:1000 dilution. GSDMD was detected using anti-GSDMD (Abcam, Cat#ab219800) at 1:1000 dilution. NINJ1 was detected using anti-NINJ1 (kindly provided by Genentech, Cat#Ninj1-rbIgG-25:10363). Tissue fibrin was detected using anti-fibrin (59D8) at 1 mg/ml. Images were acquired with LI-COR Odyssey Imager.

### Isolation of MVs from mouse plasma

Plasma was collected as mentioned above in PT. Then 50 μL of mouse plasma was diluted with 1 mL of HBSA (137 mM NaCl, 5.38 mM KCl, 5.55 mM glucose, 10 mM HEPES, 0.1% bovine serum albumin, pH 7.5). MVs were pelleted at 20,000 g for 60 minutes at 4 °C, washed once with 1 mL of HBSA and resuspended in 100 μL HBSA.

### Isolation of MVs from cell culture medium

Cell debris was removed from cell culture supernatant by centrifugation at 1,000 g for 10 minutes. MVs were then pelleted at 20,000 g for 60 minutes at 4 °C and resuspended in HBSA.

### MV Counting

To count MVs collected from cell culture supernatant, cell debris was removed by centrifugation at 1000 g for 10 minutes. MVs in cell culture supernatant or plasma were analyzed directly without further centrifugation. All samples were diluted to optimal conditions (about 10^8^ particles/mL) for analysis in PBS. Video recordings were made for 10 intervals at a length of 30 seconds each using NanoSight. Nanoparticle tracking analysis was performed to measure the number and size of the plasma and culture supernatant MVs.

### Plasma MV TF Activity Assay

Plasma samples (25 μL each) were incubated with the 1H1 anti-TF antibody (Genentech) at 100 mg/ml or rat IgG controls for 15 minutes at room temperature. Next, 25 μL of HBSA containing 10 nM mouse FVIIa, 300 nM human FX and 10 mM CaCl2 was added to the samples and incubated for 2 hours at 37 °C in a half area 96-well plate. Finally, 12.5 μL of the FXa substrate RGR-XaChrom (4 mM, Enzyme Research Laboratories#100-03) was added and the mixture was incubated at 37 °C for 15 minutes. Absorbance at 405 nm was measured on Cytation 5. The relative TF activity was calculated with absorbance value after subtracting the TF-independent activity in the presence of TF blocking antibodies from the total activity in the presence of the IgG controls.

### Statistical analysis

Data are represented as individual dots, with bar denoting mean. For multiple-group with two independent factors, two-way ANOVA with Holm-Sidak multiple comparisons was used for normally distributed variables. A p value <0.05 was considered significant. Statistical analyses were performed in Prism GraphPad Prism 9.

## Acknowledgements

This work was supported by the National Institutes of Health R00 HL145117 to C.W., R01 HL142640 and GM132443 to Y.W. and Z. L., R01 HL146744 to Z.L., and HL129193 to J.P.W. Dr. Wendy Katz provided help with tissue paraffin embedding and sectioning and was supported by NIH/NIGMS Institutional Development Award P20GM103527. Dr. Hartmut Weiler at Medical College of Wisconsin and Dr. Rodney M. Camire at the University of Pennsylvania provided fibrin antibody. *Ninj1^+/−^* mice and NINJ1 antibody were provided by Genentech Inc. Dr. Chris Richard provides help with MV counting on Nanosight.

## Author contributions

C.W. and J.C. designed and performed the experiments and wrote the manuscript. H.L., G.Z., Y.Z., L.Y., and M.M.S.S. assisted the experiments. J.P.W., Y.W, and Z.L. contributed to manuscript preparation. All authors discussed the results and commented on the manuscript.

## Corresponding authors

Correspondence and requests for materials should be addressed to.

## Supplementary Data

**Figure S1.**
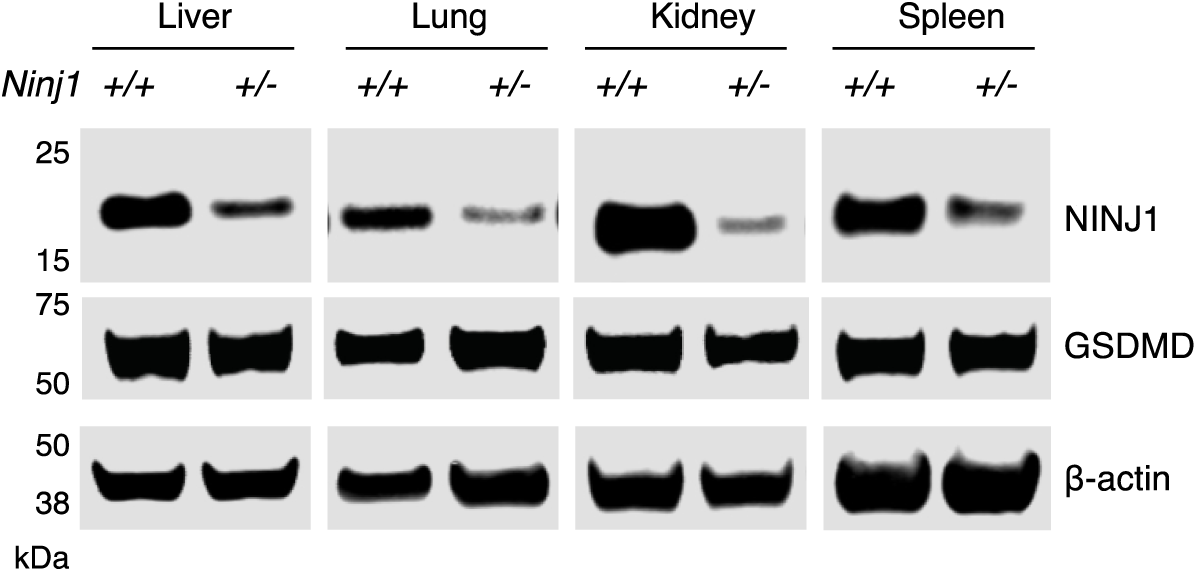
NINJ1 protein abundance in different tissues. Protein was extracted from fresh frozen tissues and detected by immunoblot.

**Figure S2.**
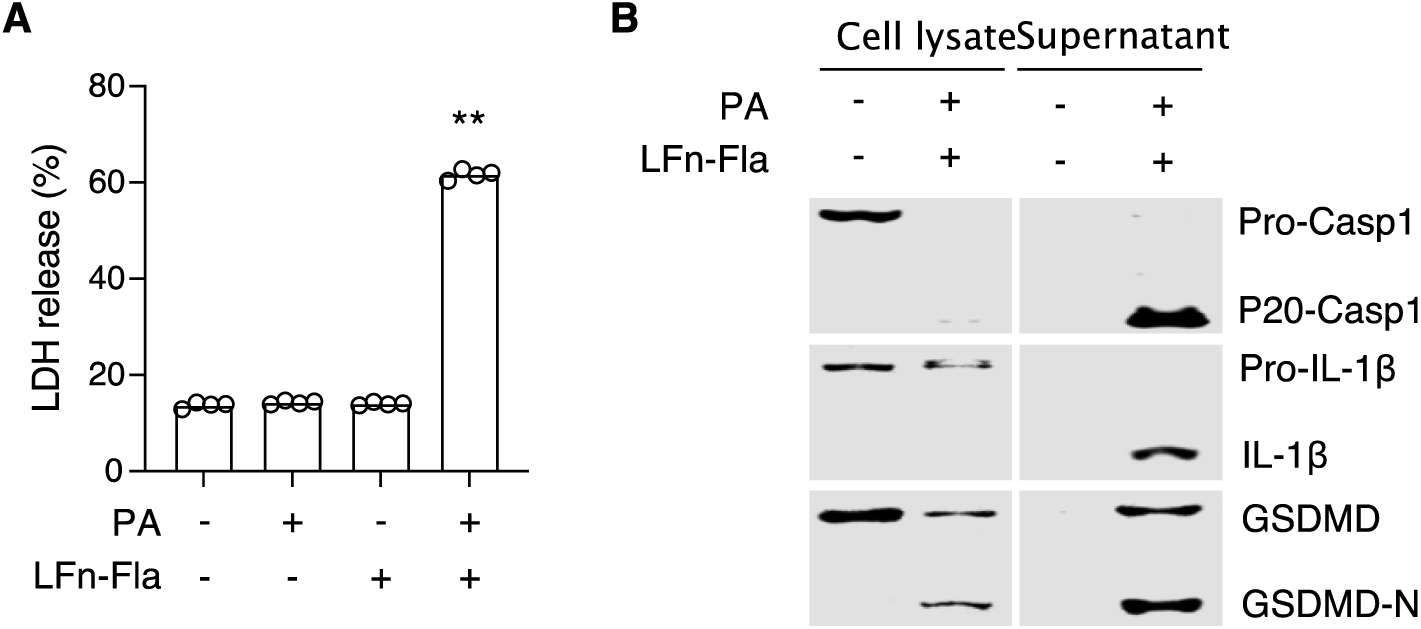
Flagellin-induced inflammasome activation and pyroptosis. BMDMs from. *Ninj1^+/+^* mice were incubated with LFn-Fla (1 μg/mL) and/or PA (1 μg /mL) for 90 minutes. (**A**) Plasma membrane rupture (PMR) was measured by LDH release. (**B**) Caspase-1, IL-1β, and GSDMD in the cell lysates and culture supernatant was detected by immunoblot. Circles represent individual mouse, with bars donating mean. ** p < 0.01 versus each of the other three groups (Student’s *t*-test; unpaired).

**Figure S3.**
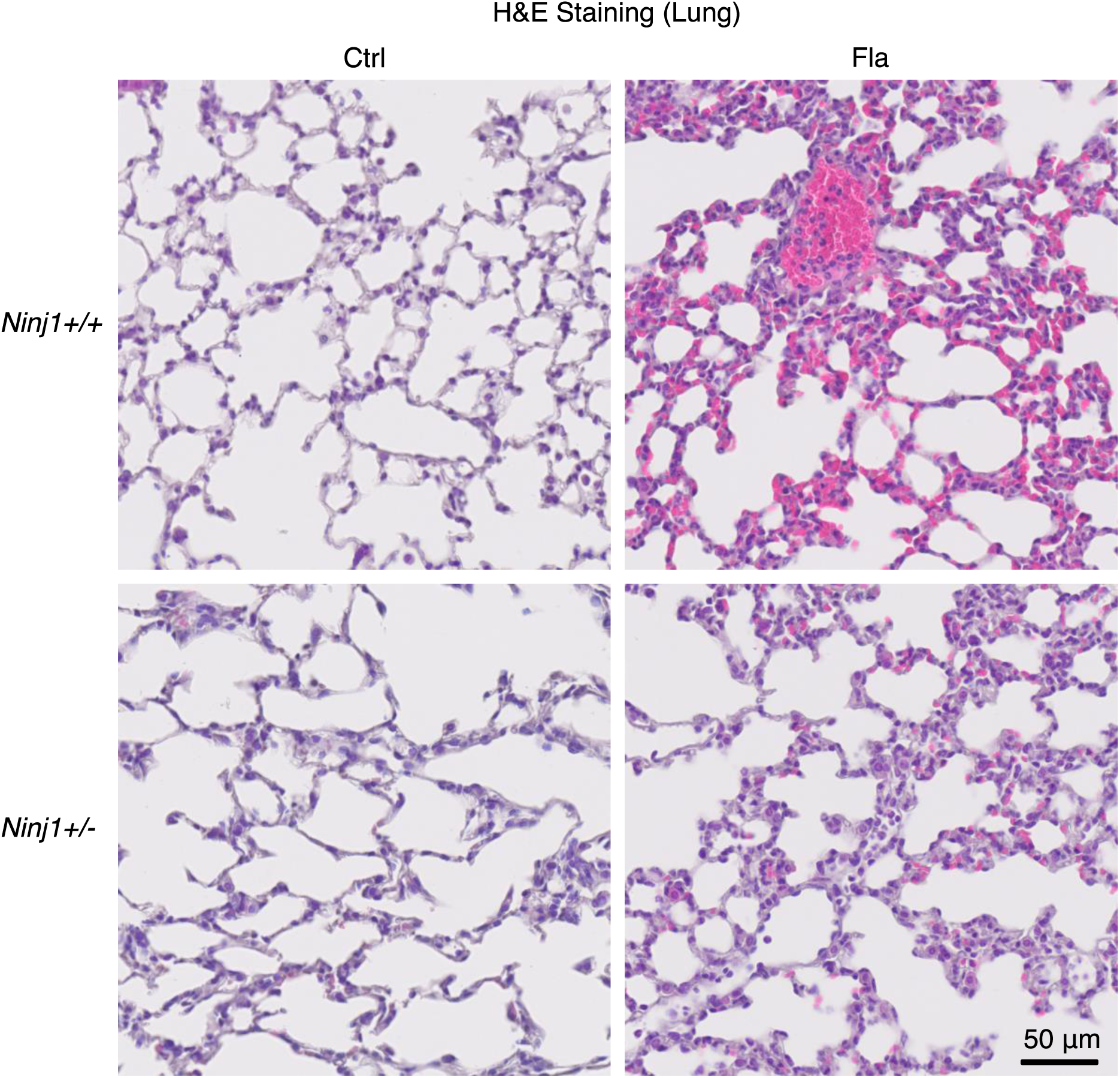
Flagellin-induced pathology in *Ninj1^+/+^* and *Ninj1^+/−^*. Mice were injected intravenously with Ctrl (saline) or Fla (500 ng LFn-Fla plus 3 μg PA). Tissues was collected 90 minutes after Ctrl or Fla injection. H&E staining was performed on saline-perfused and paraffin-embedded lung tissue sections.

**Figure S4.**
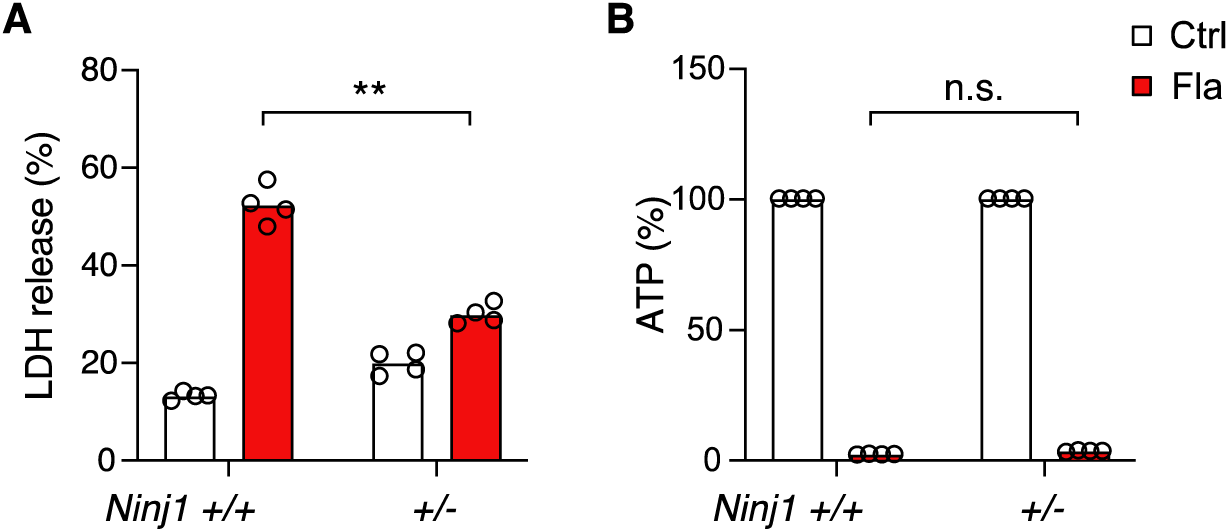
Flagellin-induced pyroptosis in *Ninj1^+/+^* and *Ninj1^+/−^* BMDMs. BMDMs from *Ninj1^+/+^*and *Ninj1^+/−^* mice were incubated with Ctrl (PBS) or Fla (1 μg/mL LFn-Fla plus 1 μg/mL PA) for 90 minutes. LDH release (**A**) and ATP (**B**) were measured. Circles represent individual mouse, with bars donating mean. ** p < 0.01, n.s. donates not significant (two-way ANOVA with Holm-Sidak multiple comparisons).

**Figure S5.**
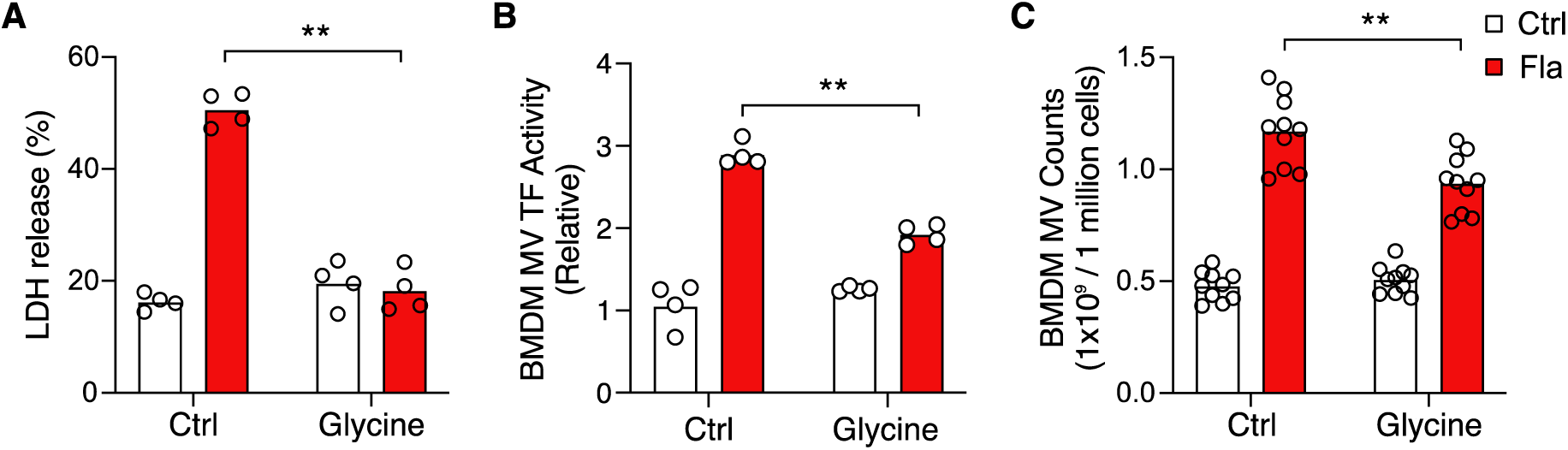
Glycine treatment inhibits BMDM MV release. (**A-C**) BMDMs from *Ninj1^+/+^* mice were treated with 5 mM glycine for 30 minutes before incubation with Ctrl (PBS) or Fla (1 μg/mL LFn-Fla plus 1 μg/mL PA). Cell culture supernatant was collected after 90 minutes incubation. LDH release (**A**), BMDM MV TF activity (**B**), and MV counts (**C**) were measured. Circles represent individual mouse, with bars donating mean. ** p < 0.01, n.s. donates not significant (two-way ANOVA with Holm-Sidak multiple comparisons).

