## Supplementary material for "Inhibiting NINJ1-dependent plasma membrane rupture protects against inflammasome-induced blood coagulation and inflammation": suppl

### Supplementary Data

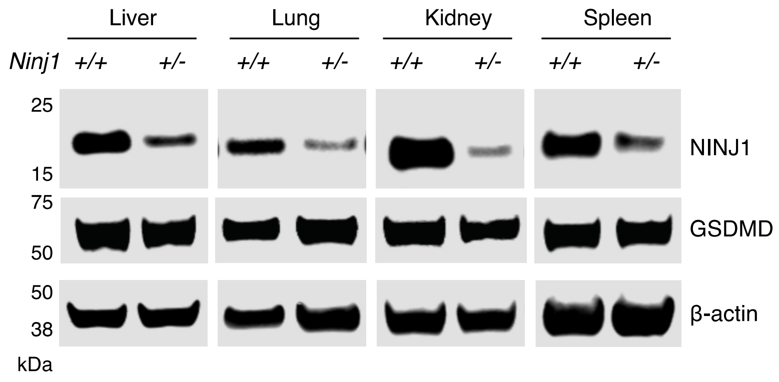


**Figure S1**. **NINJ1 protein abundance in different tissues.** Protein was extracted from fresh frozen tissues and detected by immunoblot.

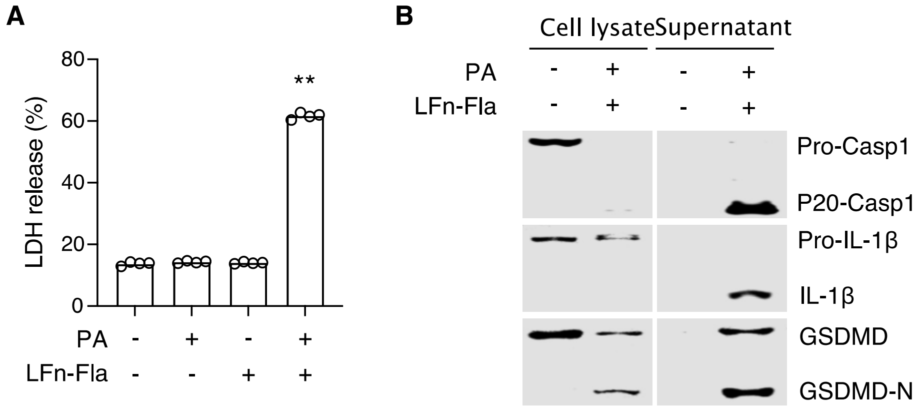


**Figure S2**. **Flagellin-induced inflammasome activation and pyroptosis.** BMDMs from *Ninj1*$^{+/+}$ mice were incubated with LFn-Fla (1 μg/mL) and/or PA (1 μg /mL) for 90 minutes. (**A**) Plasma membrane rupture (PMR) was measured by LDH release. (**B**) Caspase-1, IL-1β, and GSDMD in the cell lysates and culture supernatant was detected by immunoblot. Circles represent individual mouse, with bars donating mean. ** $p < 0.01$ versus each of the other three groups (Student's *t*-test; unpaired).

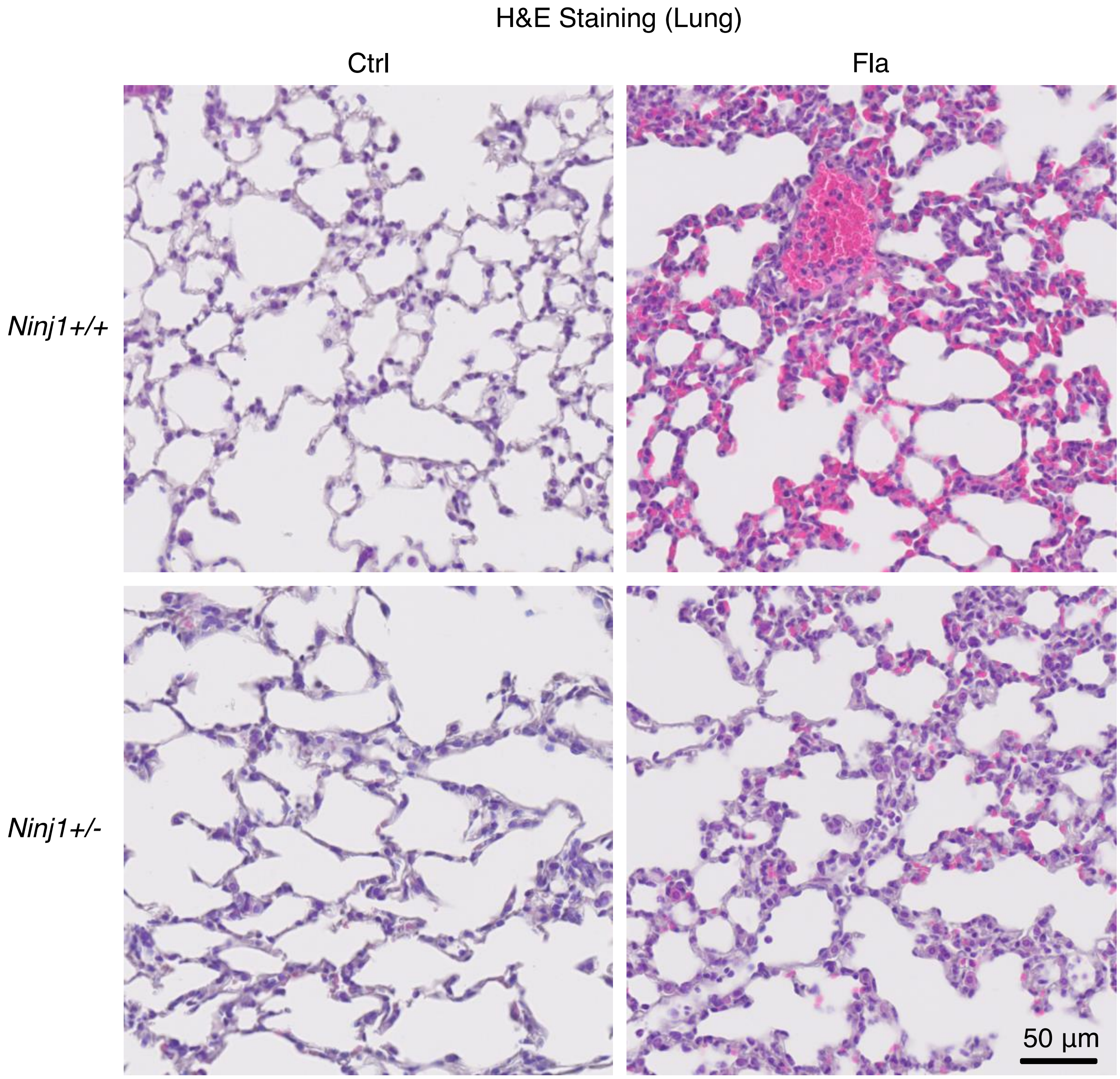


**Figure S3. Flagellin-induced pathology in *Ninj1*$^{+/+}$ and *Ninj1*$^{+/-}$.**

Mice were injected intravenously with Ctrl (saline) or Fla (500 ng LFn-Fla plus 3 µg PA). Tissues was collected 90 minutes after Ctrl or Fla injection. H&E staining was performed on saline-perfused and paraffin-embedded lung tissue sections.

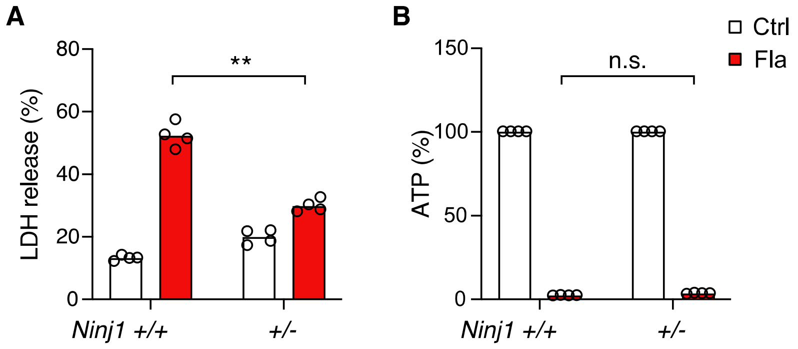


**Figure S4. Flagellin-induced pyroptosis in *Ninj1*$^{+/+}$ and *Ninj1*$^{+/-}$ BMDMs**. BMDMs from *Ninj1*$^{+/+}$ and *Ninj1*$^{+/-}$ mice were incubated with Ctrl (PBS) or Fla (1 µg/mL LFn-Fla plus 1 µg/mL PA) for 90 minutes. LDH release (**A**) and ATP (**B**) were measured. Circles represent individual mouse, with bars donating mean. ** $p < 0.01$, n.s. donates not significant (two-way ANOVA with Holm-Sidak multiple comparisons).

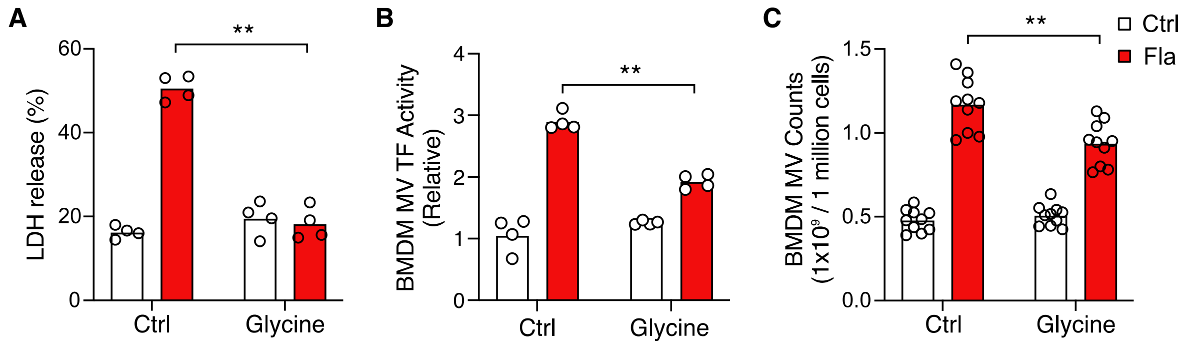


**Figure S5. Glycine treatment inhibits BMDM MV release.** (**A-C**) BMDMs from *Ninj1*$^{+/+}$ mice were treated with 5 mM glycine for 30 minutes before incubation with Ctrl (PBS) or Fla (1 μg/mL LFn-Fla plus 1 μg/mL PA). Cell culture supernatant was collected after 90 minutes incubation. LDH release (**A**), BMDM MV TF activity (**B**), and MV counts (**C**) were measured. Circles represent individual mouse, with bars donating mean. ** $p < 0.01$, n.s. donates not significant (two-way ANOVA with Holm-Sidak multiple comparisons).
